# Gut Microbiome Alterations in Canine Idiopathic Epilepsy: A Pairwise Case-Control Study

**DOI:** 10.64898/2026.04.02.716098

**Authors:** Yixuan Yang, Julie Nettifee, M. Andrea Azcarate-Peril, Karen R. Muñana, Benjamin Callahan

## Abstract

**Background:** Idiopathic epilepsy (IE) is the most common chronic nervous system disorder of dogs, and its cause is poorly understood. Emerging evidence suggests that microbiome alterations can occur with IE via the microbiota-gut-brain axis. Therefore, we analyzed the fecal microbiomes of 98 dogs (49 IE, 49 control) in a pairwise case-control observational study using 16S rRNA gene sequencing.

**Results:** Although the microbial community was mostly similar between groups, IE was associated with a modest but significant shift in Weighted-Unifrac distance (P = 0.042). We used six differential abundance (DA) methods to identify differentially abundant amplicon sequencing variants (ASVs) between IE and control groups. Notably, one *Collinsella* ASV was found to be significantly more abundant in IE dogs by all six methods. The gut microbial compositions varied drastically across households (accounting for about 69% of the total variation), but did not have significant differences between sex, age, or breed. Phenobarbital administration in IE dogs had a significant effect on seizure control, and was not associated with changes in the microbiome.

**Conclusion:** Our findings suggest a relationship between gut microbiomes and IE. However, the specific mechanism needs to be further investigated.

## Background

Epilepsy is the most common chronic neurological disorder of dogs[1], with an estimated prevalence of 0.62-0.75% in the general population[2–4]. Over half of all dogs with epilepsy are diagnosed with idiopathic epilepsy (IE), a clinical syndrome characterized by recurrent seizures for which there is no underlying cause other than a presumed genetic predisposition[5–7]. Despite its prevalence, IE remains a poorly understood disorder, for which the standard of care is limited to symptomatic therapy with antiseizure medications (ASMs). Furthermore, medical management is ineffective in controlling seizures in approximately one-third of dogs with IE[8], and these dogs suffer an increased morbidity and fatality associated with their disease[5, 9].

The microbiota-gut-brain axis, a complex connection between the gastrointestinal tract and the nervous system, has been implicated in the development and progression of neurological disorders of humans, including multiple sclerosis[10, 11], Parkinson’s disease[12–14], Alzheimers disease[15, 16] and autism spectrum disorder[17, 18].

Evidence for a similar association between epilepsy and the gut microbiota in humans is growing. A recent meta-analysis that included 12 case control studies involving 807 participants identified significant differences in gut microbial composition between epileptic patients and healthy controls, with an overall reduction in gut microbial diversity[19]. Several recent publications suggest that alterations in the gut microbiome might also be present in dogs with epilepsy. A study comparing the fecal microbiota of drug-naïve idiopathic epileptic dogs and healthy controls identified epileptic dogs to have a decrease in bacterial richness, with alterations in the relative abundance of several bacterial genera[20]. A second study evaluated 49 dogs with epilepsy and 39 healthy controls identified alterations in both the fecal metabolome and microbial composition in dogs with epilepsy, with additional differences noted among epileptic dogs categorized as being drug resistant or having a mild phenotype[21]. Our lab previously performed a pilot study comparing the fecal microbiome in dogs with IE to healthy housemates with a specific focus on Lactobacillus, in which a strong household effect was identified[22]. The current study aims to build on this work, utilizing a paired household design, while strictly controlling for the strongest confounders (household/diet/environment) to test the following hypotheses: (1) Dogs with IE possess reproducible gut microbiota signatures that extend beyond the household effect; 2) Changes in the abundance of specific microbial taxa are independently associated with IE status; and 3) Among dogs with IE, specific microbial patterns are associated with seizure frequency and response to treatment. Since microbiome differential abundance (DA) results can vary drastically depending on the statistical framework used, we followed recent recommendations to adopt a consensus approach[23]. We employed an ensemble of six tools to ensure that our findings are robust and independent of specific tool assumptions.

## Results

### Study Population

Survey information and fecal samples were obtained from 118 dogs residing in 51 households with at least one IE and one control dog, in a continuation of our pilot study[22]. Fecal microbiome composition was measured by 16S rRNA gene sequencing. We studied a subset of the enrolled cohort, 98 dogs representing 49 pairs of IE and control dogs from 49 households. Two households were excluded because of insufficient sequencing depth from a fecal sample, and for households that contributed >2 dogs, only one IE and one control animal were used (see *Methods*). Dogs in the same household were fed the same diet. These households are distributed among 23 states in the U.S., with the largest number (n=8) from North Carolina. Dog breed was reported by the owner and breeds were grouped into seven breed categories based on genetic similarities as described by Dutrow et al.[24]: herder, pointer spaniel, retriever, terrier, scent hound, sight hound, and sled dog. Dogs that shared similarities with more than one breed category or shared characteristics with most of the breed groupings, and dogs of breeds that were not included in the Dutrow study were excluded from breed analysis. In *Table 1,* a summary of the study cohort is presented. Breed categories were evenly distributed between the IE and control group (Chi-Square, p = 0.9887). The ages of the dogs are evenly distributed between the IE and control groups with a mean of approximately 75.6 months (t-test, p=0.9955). Fifty-eight of the 98 dogs included in our study were female. A higher IE rate was found in male dogs (60%) than female dogs (43.1%) in line with previous reports[25–27]. However, this difference did not reach the standard level of statistical significance in this study alone (Chi-Square, p=0.15). Phenobarbital had a significant effect on seizure freedom compared to no treatment (Chi-Square, p = 0.009). Among the 19 IE dogs that were receiving phenobarbital, 11 (58%) were seizure free; among the 29 drug-naive dogs, only 5 were noted to be free from seizures.

**Table 1:**
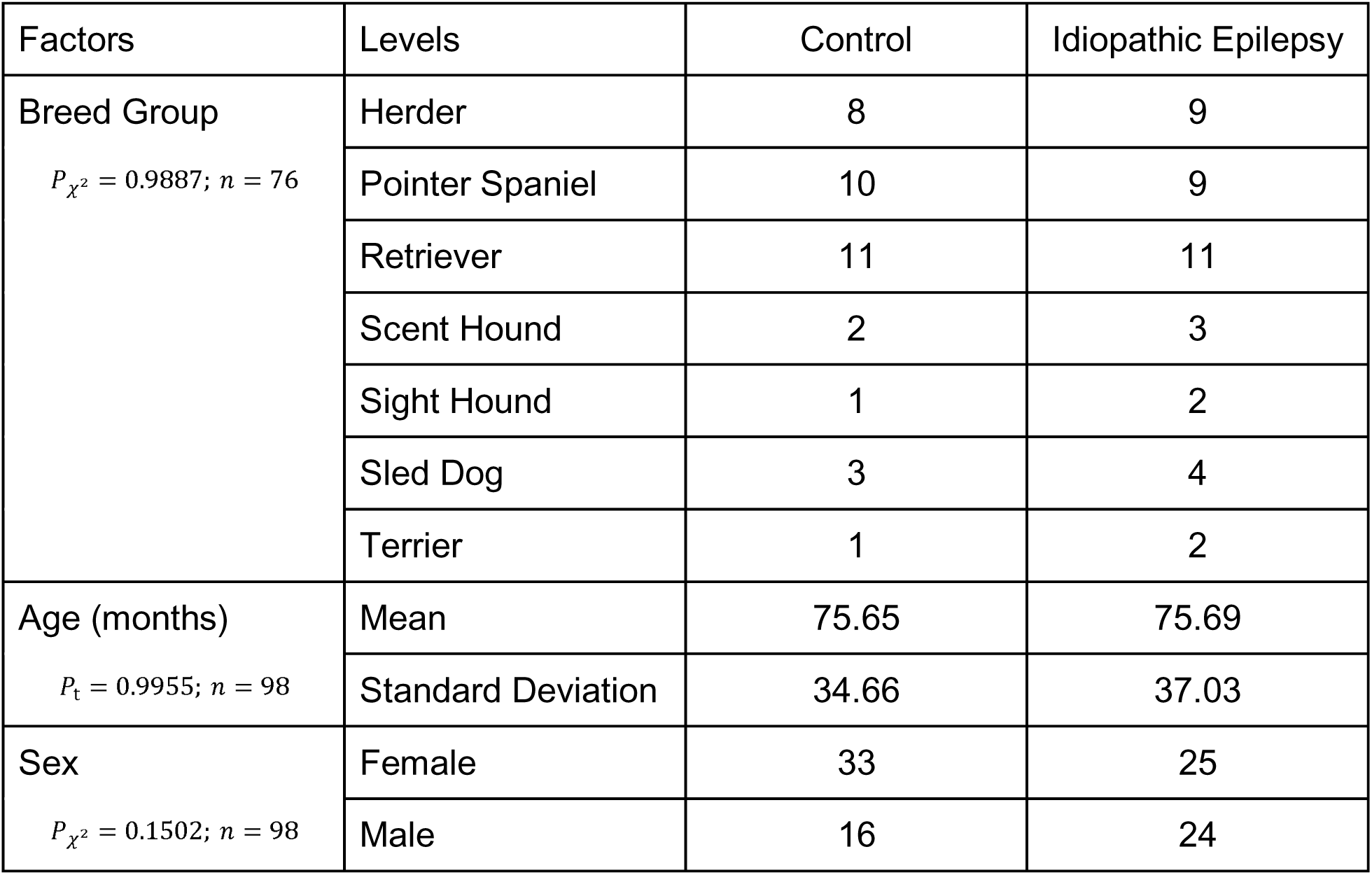
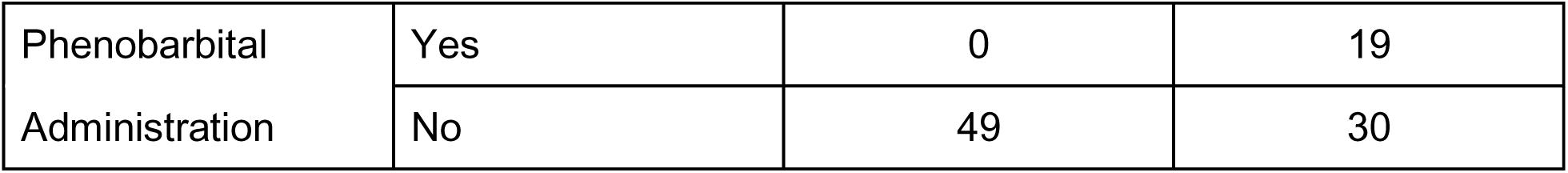
Summary information of the study cohort (N=98). P-values are reported for comparisons of breed group, age and sex distributions between the paired IE (N=49) and control (N=49) subgroups. Only a subset of N=76 dogs with defined breed groups are reported in the breed group section.

### Fecal Microbiome Composition Measured in this Study was similar to that measured in our Pilot Study

As the present study is a continuation of the pilot study, the microbial composition was compared between these two studies. The distribution of bacterial relative abundances at the phylum level was consistent between the pilot study and this study (Figure 1). In the pilot and the present study, respectively, 75.3% and 76.1% of amplicon sequence variants (ASVs) were classified as Firmicutes, 10% and 12.2% of ASVs were classified as Actinobacteriota, 9.2% and 5.7% of ASVs were classified as Bacteroidota on average. Unlike other phyla, the abundance of Actinobacteria in our study was almost entirely attributed to a single genus: *Collinsella* made up a large majority of Actinobacteriota in both studies (pilot study: 87.6% ± 7.5%; present study: 80.3% ± 15.6%).

**Figure 1:**
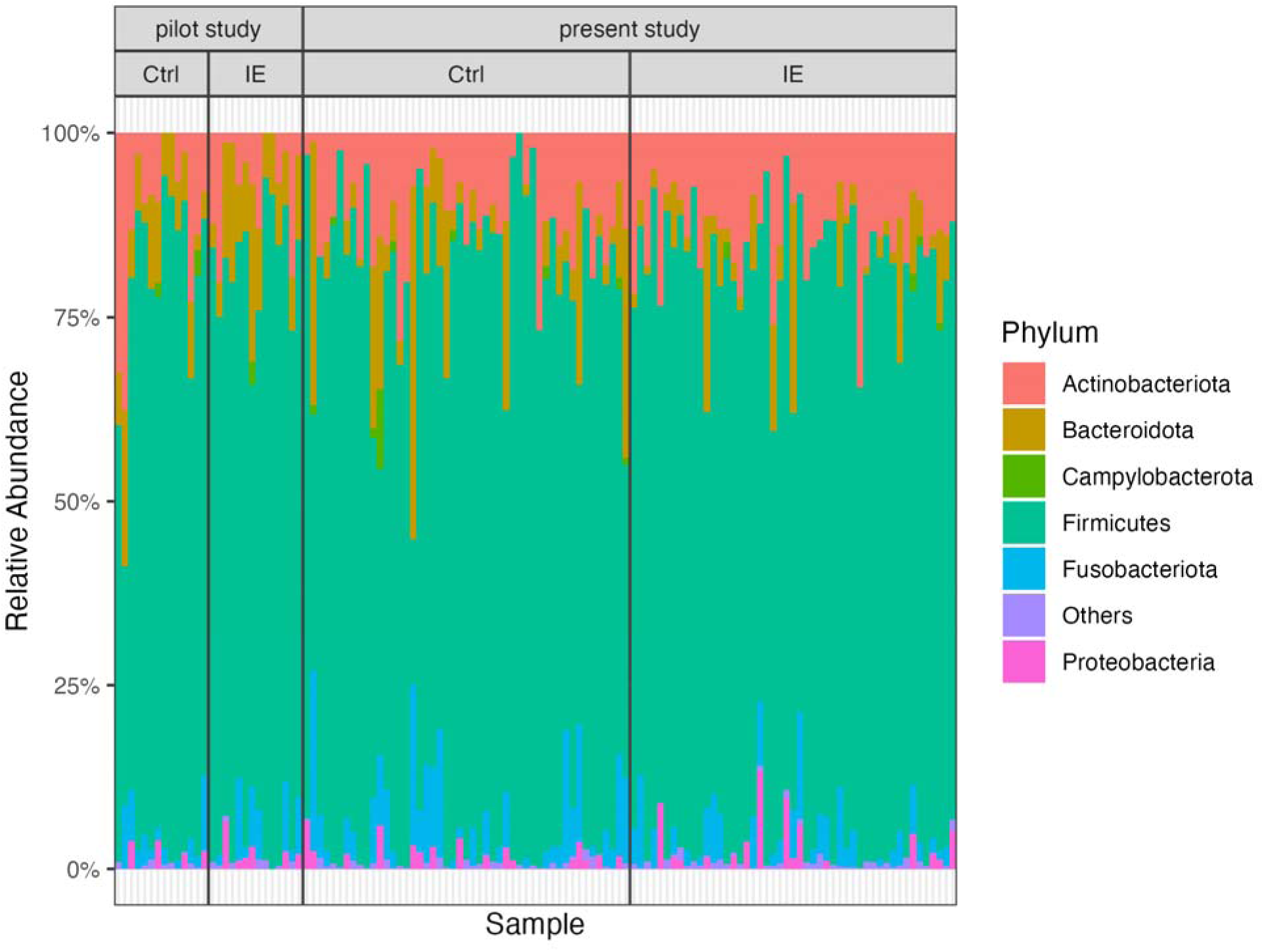
Relative abundances of bacterial phyla in fecal samples from the pilot and this study. Phyla with relative abundance less than 1% were classified as Others.

### Household Explains Most of the Microbial Community Variation

In the previous pilot study, household was reported as the most influential factor associated with the composition of the fecal microbial community at the ASV level. That is, dogs from the same household had much more similar gut microbiomes than those from different households. In this study, household was again found to be the strongest factor explaining gut microbiome composition based on analysis of community-wise distances (PERMANOVA, Bray-Curtis, p-value < 0.0001; PERMANOVA, Weighted-Unifrac, p-value < 0.0001). Furthermore, household was also estimated to explain most of the variation of the gut microbiome (PERMANOVA, Bray-Curtis, R^2^ = 0.689; PERMANOVA, Weighted-Unifrac, R^2^ = 0.677). Breed group, age, sex and phenobarbital administration were not significantly associated with microbiome composition (Table 2). The difference in microbiome composition between IE and control dogs was marginally significant (PERMANOVA, Bray-Curtis, p-value = 0.090; PERMANOVA, Weighted-Unifrac, p-value = 0.042). The household effect was controlled for by leveraging the paired study design in all of these tests (see *Methods*).

**Table 2:** P-values and effect sizes for comparisons of fecal microbiome composition at ASV level across several different factors. Permutational Multivariate Analysis of Variance (PERMANOVA) analysis was performed on both Bray-Curtis and Weighted-Unifrac community dissimilarity metrics. ANOVA analysis of alpha-diversity used the Shannon diversity index. When testing the breed group factor, only samples with a defined breed group were included (N = 76). When testing the phenobarbital administration factor, only samples in the IE group were included (N=49).

| Methods Factors | p-value | | | $R^2$ | | |
| --- | --- | --- | --- | --- | --- | --- |
|  | PERMANOVA (Bray-Curtis) | PERMANOVA (Weighted-Unifrac) | ANOVA (Shannon Index) | PERMANOVA (Bray-Curtis) | PERMANOVA (Weighted-Unifrac) | ANOVA (Shannon Index) |
| Household | **<0.0001** | **<0.0001** | 0.053 | 0.689 | 0.678 | 0.610 |
| IE status | 0.090 | **0.042** | 0.796 | 0.009 | 0.014 | 0.001 |
| Breed group | 0.143 | 0.101 | 0.770 | 0.049 | 0.058 | 0.078 |
| Age | 0.523 | 0.490 | 0.920 | 0.006 | 0.006 | 0.0002 |
| Sex | 0.981 | 0.997 | 0.155 | 0.003 | 0.001 | 0.042 |
| Phenobarbital Administration | 0.437 | 0.743 | 0.343 | 0.021 | 0.014 | 0.019 |

### Dogs from the Same Household Share Bacterial ASVs

To better understand the household effect, we examined the fraction of bacterial ASVs shared between pairs of dogs from the same household. ASVs represent unique variants of the V4 rRNA gene sequenced in this study, and are the finest level of taxonomic resolution possible from such data. We first calculated the Jaccard similarity – the fraction of ASVs shared between two samples divided by the total number of ASVs detected in either sample – between the paired IE/control dogs from each of our 49 households. Then, we created a new dataset in which each IE dog was paired with a randomly selected control dog from a different household while forcing the predominant breed to be the same as possible to control the breed effect. We observed a significantly higher overlap of ASVs between dogs from the same household than between dogs from different households (pairwise t-test, P < 0.0001), with an average of 34.7% of the ASVs detected in one dog’s fecal microbiota also detected in the other dog from their household (Figure 2).

**Figure 2:**
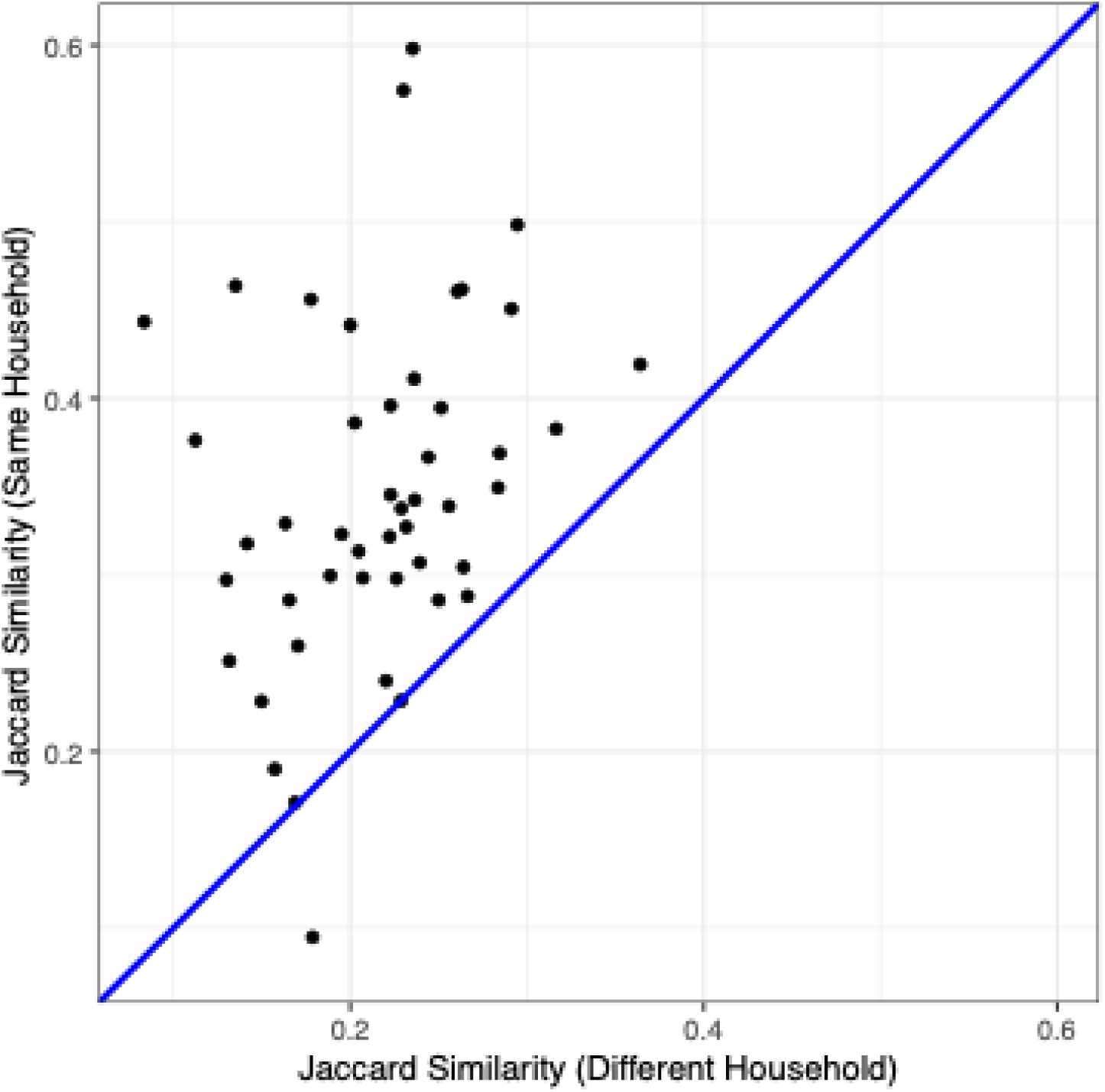
More ASVs are shared between dogs from the same household than between dogs from different households. ASV overlaps measured by Jaccard similarity between IE dogs and control dogs from a different household (x-axis) and between each IE dog and control dog from the same household (y-axis). The blue line shows the y = x expectation if the household has no effect on ASV overlap.

Next, we evaluated the frequency of sharing within a household across the ASVs detected in our study. For a random pair of dogs, the expected probability of one ASV present in both dogs is the square of the prevalence for that ASV. Most ASVs — 286 out of 322 ASVs with prevalence greater than 0.1 — were shared within a household more often than expected by their study-wide prevalence (Figure 3), suggesting the household effect acts broadly across the many taxa present in the canine fecal microbiome.

**Figure 3:**
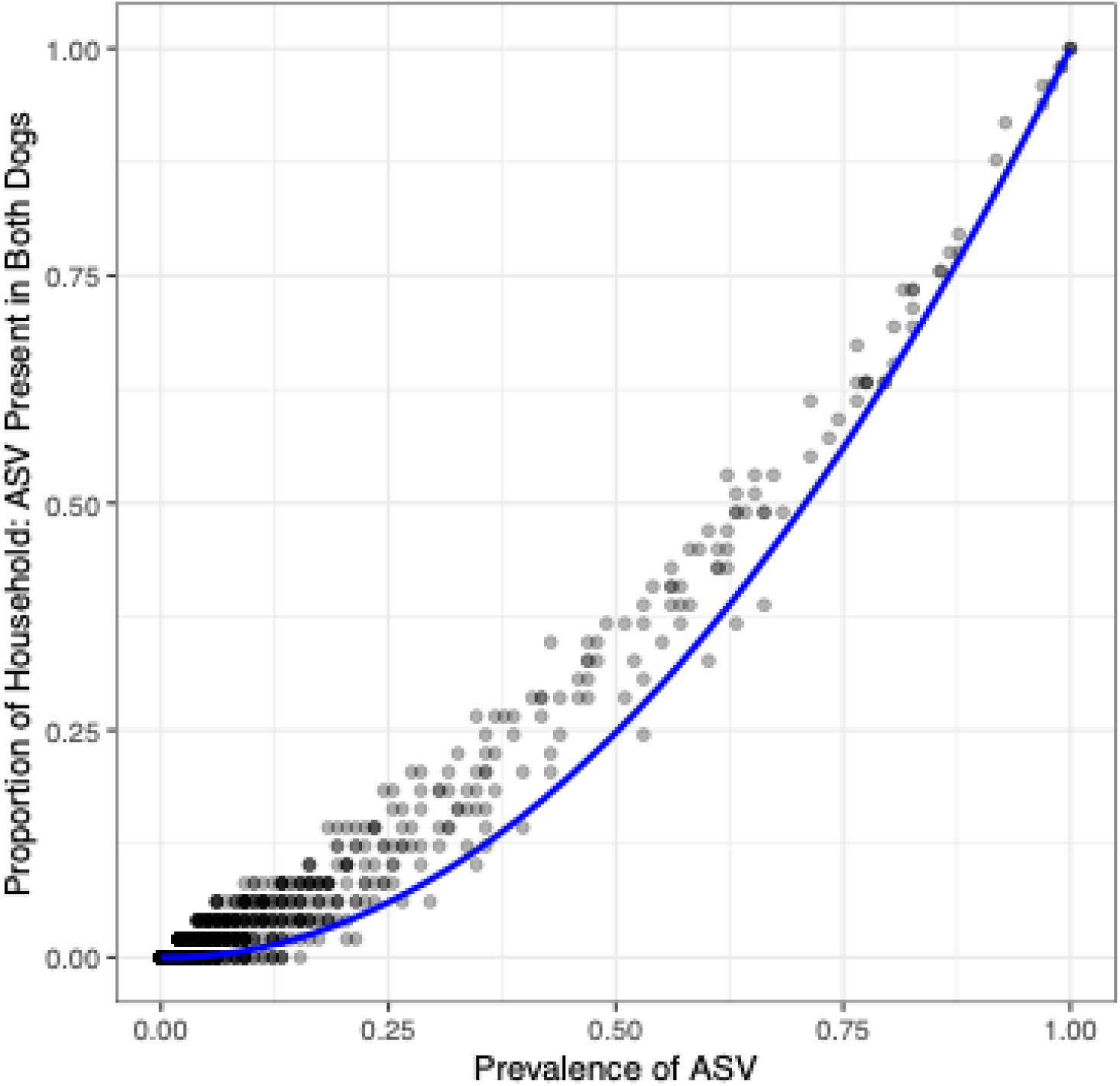
The proportion of households in which an ASV was present in both the IE and control dogs as a function of study-wide ASV prevalence. The y-axis represents the observed proportion of households in which the ASV was present in both dogs. The x-axis represents the study-wide prevalence of each ASV. Each point represents an ASV present in our study. The blue curved line represents the expected proportion of households for which the ASV would occur in both dogs if dogs were paired randomly, adjusted for sampling without replacement within the finite study population (n=98).

### Differential Abundance of ASVs Between IE and Control Dogs

We investigated whether ASVs were differentially abundant (DA) between IE and control dogs using a consensus approach incorporating multiple common DA tools including pairwise t-test-based tools such as ALDEx2 t-test and R-base t-test, and linear regression-based tools such as ALDEx2 GLM, ANCOM-BC, corncob, and LinDA[28–31]. The household effect was controlled for by leveraging the paired study design in all of these tests (see *Methods*). To improve statistical power, ASVs with prevalence lower than 10% were excluded.

The identification of significantly differentially abundant ASVs was generally consistent across DA methods, with one *Collinsella* ASV consistently identified by all six tools (Table 3). ALDEx t-test, ALDEx GLM, ANCOM-BC, and linDA showed high correlations in p-values between each other, while corncob and a paired t-test on rarefied count data produced p-values that were correlated but less strongly (Figure 4a). The estimated log2 fold changes (LFC) from different tools for consensus significant ASVs were all consistent in their direction of effect (Figure 4b).

**Figure 4:**
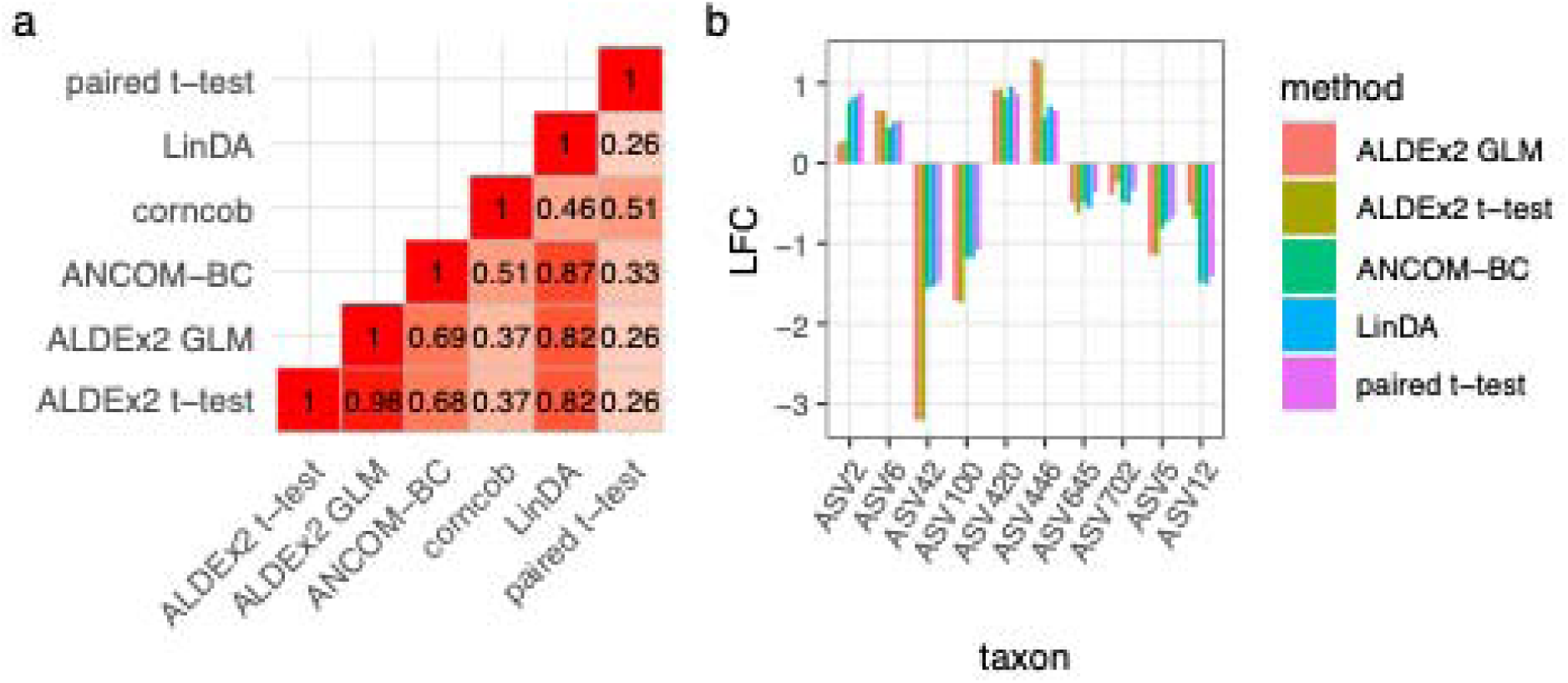
The consistency of differential abundance (DA) results at the ASV level across six different DA methods. (a) The correlation between p-values returned from each DA tool considered. (b) The log-fold change (LFC) effect size of the top 10 ASVs that were reported as significantly differential in abundance between IE and control dogs. Corncob was not included since the method does not calculate LFC. However, the direction of relative abundance difference estimated by corncob agreed with the other DA methods for all of these ASVs.

**Table 3:** Top 10 candidate differentially abundant (DA) ASVs based on consensus reporting by six different DA tools. Genus-level taxonomic assignments that were not identified by DADA2 taxonomy assignment were filled in with the top alignment of the BLAST software against the nt/nr database. The genus of ASV645 remained ambiguous.

| Taxon | Phylum | Genus | ALDEx2 t test | ALDEx2 GLM | ANCOM-BC | corncob | LinDA | paired t test |
| --- | --- | --- | --- | --- | --- | --- | --- | --- |
| ASV2 | Actinobacteriota | *Collinsella* | **0.0056** | **0.0062** | **0.0004** | **0.0003** | **0.0094** | **0.0109** |
| ASV6 | Firmicutes | *[Ruminococcus] gnavus* group | **0.0277** | **0.0276** | **0.0139** | **0.0088** | **0.0466** | 0.1792 |
| ASV42 | Firmicutes | *[Ruminococcus] torques* group | 0.0930 | 0.0905 | **0.0002** | **0.0404** | **0.0162** | **0.0398** |
| ASV100 | Firmicutes | *Lachnoclostridium* | 0.1793 | 0.1827 | **0.0001** | **0.0001** | **0.0148** | **0.0111** |
| ASV420 | Firmicutes | *Clostridium sensu stricto 1* | 0.1560 | 0.1755 | **0.0009** | **0.0001** | **0.0178** | **0.0288** |
| ASV446 | Firmicutes | *Mediterraneibacter* | 0.1238 | 0.1128 | **0.0007** | **0.0002** | **0.0124** | **0.0340** |
| ASV645 | Firmicutes | NA | 0.4058 | 0.3948 | **0.0011** | **0.0002** | **0.0193** | **0.0292** |
| ASV702 | Firmicutes | *Holdemania* | 0.4791 | 0.4769 | **0.0002** | **0.0000** | **0.0111** | **0.0209** |
| ASV5 | Fusobacteriota | *Fusobacterium* | 0.2526 | 0.2733 | **0.0206** | **0.0101** | 0.1318 | **0.0310** |
| ASV12 | Firmicutes | *Blautia* | 0.1137 | 0.1055 | **0.0003** | **0.0355** | **0.0172** | 0.1086 |

**Figure 5:**
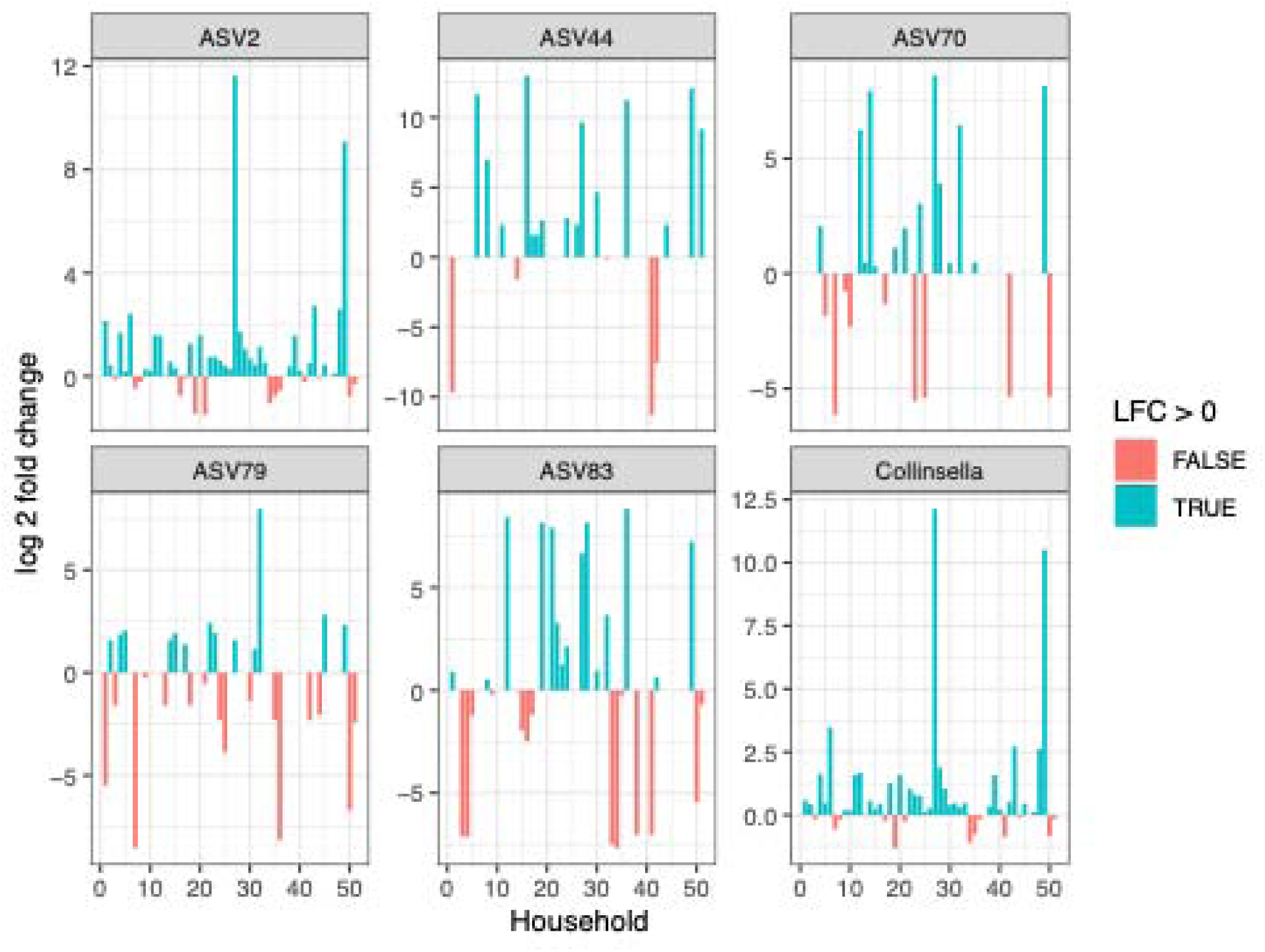
The log2 fold difference in relative abundance between IE and control dogs from the same household for the five most abundant *Collinsella* ASVs and total *Collinsella*. Absent bars indicate that the *Collinsella* ASV was not detected in either dog in that household.

### Higher Abundance of *Collinsella* Variants in IE Dogs

An ASV belonging to the genus *Collinsella* was the only ASV identified as differentially abundant by all six DA methods we considered (Table 3). *Collinsella* has also been found to be associated with neurological diseases in previous studies[16, 32, 33].

Therefore, we conducted a more in-depth investigation of *Collinsella* in our study. We considered five *Collinsella* ASVs with a prevalence higher than 10% in our study, which were assigned to four different species (Table 4). The overall prevalence of *Collinsella* was high: 97 out of 98 canine fecal samples had at least one *Collinsella* ASV. Both of the most abundant *Collinsella* ASVs – ASV2 assigned to *C. stercoris* and ASV44 assigned to *C. aerofasciens* – were significantly more abundant in the IE group by the pairwise sign test (P < 0.05). The presence of ASV79 was found to be significantly associated with sex (Chi-square, P = 0.016). The abundance of ASV83 was significantly higher in older dogs (Linear Model Regression, P = 0.002). Within IE dogs, none of the *Collinsella* ASVs were found to be significantly differentially abundant between seizure free and non-seizure free dogs or between seizure controlled and non-seizure controlled dogs (t-test, P > 0.05).

**Table 4:**
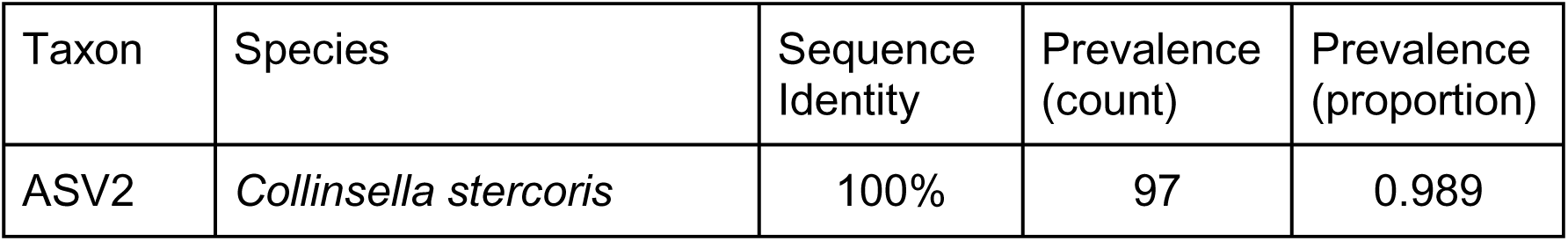
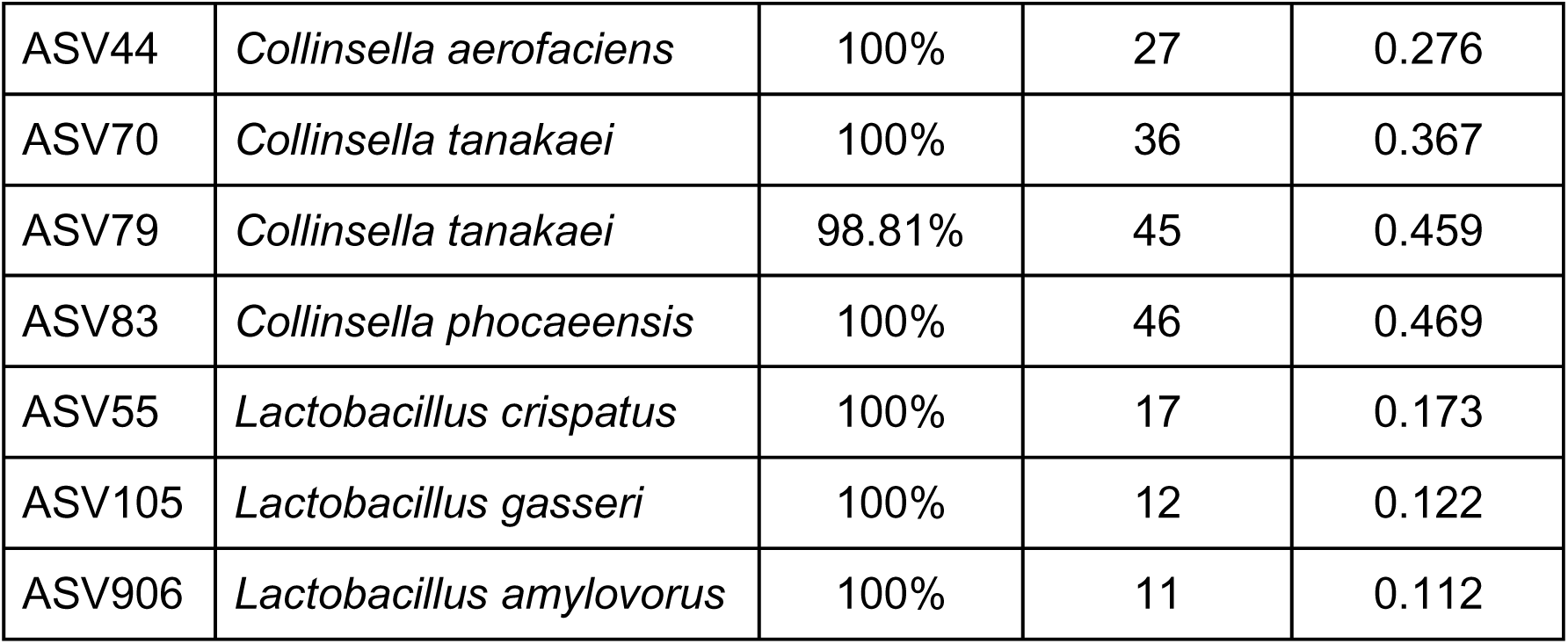
Prevalence and overall frequency of *Collinsella* and *Lactobacillus* amplicon sequence variants (ASVs) in the study population (N=98 dogs) identified through 16S rRNA gene sequencing. Species reported in this table were based on manual inspection of BLAST alignments against the nt database. Sequence identity is based on the best BLAST hit.

*Lactobacillus*, widely studied as a probiotic, has attracted growing attention for its potential beneficial role in epilepsy, with evidence emerging from both human[34] and animal[35] studies.

In the pilot study, none of the *Lactobacillus* ASVs was found to be significantly different in abundance between IE and control dogs. In this study, we identified three *Lactobacillus* ASVs with prevalences exceeding 10% (Table 4). The total *Lactobacillus* at genus level and ASV906 were found significantly more abundant in IE dogs by the pairwise sign test (P < 0.05). Sex was found marginally significant in association with ASV906 (Chi-square, P < 0.052). The abundance of total *Lactobacillus* at genus level and ASV55 was significantly higher in older dogs (Linear Regression: ASV55, P = 0.008; *Lactobacillus,* P = 0.006). ASV906 was marginally less abundant in IE dogs that are seizure controlled (t-test, P = 0.062).

### Weak Predictability of IE from the Composition of the Fecal Microbiome

As an alternative measure of the microbiome’s contribution to epilepsy, machine learning methods were applied to characterize how accurately IE can be predicted from the composition of the microbiome alone. In our balanced cohort (with equal numbers of IE and control animals), a prediction accuracy greater than 0.5 indicates the potential diagnostic value of the microbiome for IE. Random forest models, perhaps the most common type of machine learning model in microbial studies, were trained and evaluated on the ASV-level microbiome compositions. Due to the large number of ASVs and thus high dimensionality of the microbial composition, we also tried either applying

PCoA (Principal Coordinate Analysis) based on the Bray-Curtis distance or converting the data to phylum level for dimensionality reduction prior to machine learning. *Figure 6* shows the distribution of model accuracy for each dataset. The model trained by the PCoA dataset performed slightly better than the other two datasets and had a prediction accuracy of 0.588 on average. However, the model performance was not reliable enough for clinical practice.

**Figure 6:**
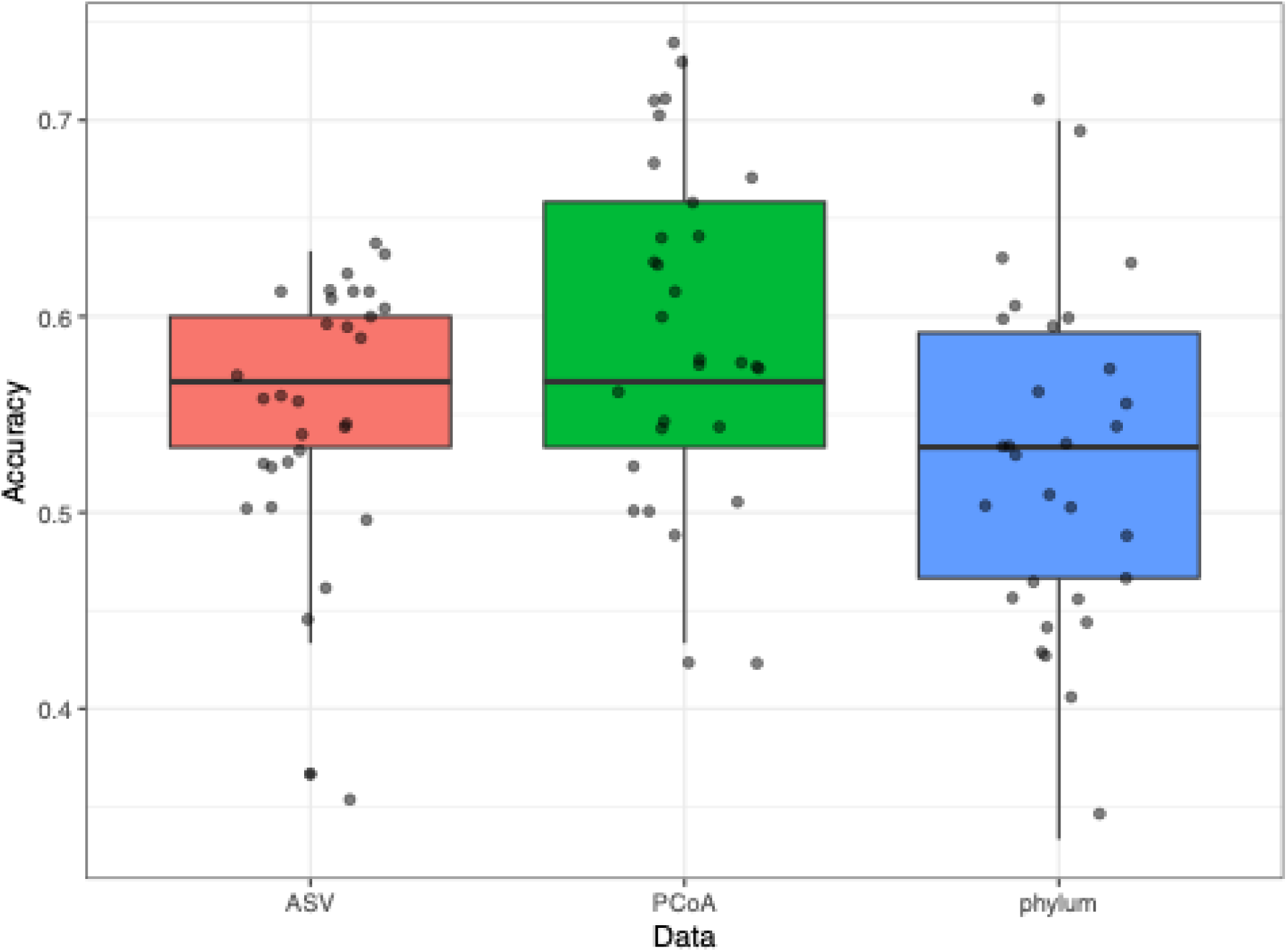
Accuracy of random forest model using three different microbiome datasets (Phylum-level, ASV-level, PCoA-level). The points represent 30 different iterations of testing and training the model. The y-axis represents simple accuracy. Median accuracy is shown by the thick horizontal line in the boxplot. Random guessing would yield 0.5 accuracy in our balanced training dataset.

## Discussion

In this study, we analyzed the fecal microbiome of 49 pairs of dogs from different households utilizing 16S rRNA sequencing, where each pair consisted of one IE dog and one control counterpart. As a continuation of the pilot study[22], we maintained a consistent study design but with a larger sample size aiming to examine the relationship between dog microbial composition and epilepsy based on the potential role of the microbiota-gut-brain axis in the development of IE. The observed phylum-level composition of fecal microbiomes in the current study was approximately the same as in the pilot study. A difference in gut microbiome between IE and control dogs was found in this larger study, and *Collinsella* was identified as the top candidate genus with relative abundance associated with IE.

In this study, household was again identified as the predominant factor influencing the canine gut microbiome, accounting for most of the observed variation. The household factor can be explained as a combination of factors, including living environment and diet, as we required that both dogs in a household had the same diet for inclusion in this study. Studies[36–38] showed that these factors can influence canine gut environments. This finding highlights the crucial role of extrinsic factors in microbial studies and should not be overlooked when interpreting the effects of other variables. Consequently, we incorporated household as a confounding factor into our subsequent analyses.

Our study design matched pairs of epileptic and non-epileptic dogs by household and diet. Previous studies have identified differences in the canine fecal microbiome based on other facotrs such as age group[39, 40], breed[41, 42], and neuter status but not sex[37, 42]. However, these studies were not designed to control for diet, one of the strongest factors influencing the gut microbiome[43], and consistent with the exceptionally strong effect of household/diet on the microbiome shown in our study .

When designing a clinical study, it is not feasible to control for the every potential confounding factors [44]. Although analysis of age and sex factors did not demonstrate statistically significant associations with microbial composition in our dataset, the lack of matching on these factors should be considered when interpreting our results. In addition, the statistical power of breed group analysis was primarily concentrated in the three largest groups (Herder, Pointer Spaniel, Retriever), while the analysis for minority groups with very few representatives remains exploratory.

Phenobarbital is considered one of the most effective treatments for canine IE. According to Watanangura et al.[45], the species diversity, microbial pattern, and taxonomic composition in the gut microbiome of dogs did not change significantly after a 90 day phenobarbital administration. Similarly, a study by Garcia-Bellenguer et al.[33] demonstrated no change in gut microbial composition of epileptic dogs following one month of treatment with phenobarbital. In this study, we also didn’t find significant differences between the fecal microbiota in IE dogs receiving phenobarbital compared to drug naive dogs, agreeing with the previous finding that phenobarbital does not alter the general pattern of the gut microbiome in dogs.

To elucidate the role of gut-microbiome in IE, the microbial taxonomy compositions were analyzed. Multiple studies have evaluated the gut microbiome difference between IE and control dogs[20–22, 33, 46]. We compared the microbiome between the IE and control dogs with differential abundance (DA) analysis at the amplicon sequence variant (ASV) level. Different DA tools rely on different statistical models and assumptions, and no single method consistently outperforms others across different datasets[23].

Therefore, we used an ensemble approach integrating the results from six DA methods to reduce false-positive findings. A *Collinsella* ASV was identified as significantly more abundant in IE dogs by every DA method applied. This concordance across multiple statistical frameworks suggests a robust association between *Collinsella* and IE, that has not been previously recognized in dogs. However, an association between *Collinsella* and epilepsy has been demonstrated in human microbiome studies[47–49]. A significant increase in the relative abundance of *Collinesella* was identified in fecal samples of children with focal epileptic seizures compared to healthy children, with the difference resolving following 3 months of treatment[49]. A subsequent study identified a significant increase in *Collinsella* in children with benign childhood epilepsy with centrotemporal spikes, a specific form of focal epilepsy, compared to healthy children[47]. Furthermore, a study of adult epilepsy patients with and without cognitive impairment identified a significant difference in relative abundance of *Collinsella* between the groups, suggesting a role for the organism in the development of cognitive dysfunction epilepsy[48]. A recent systematic review and metanalysis of the gut microbial composition in patients with epilepsy did not identify *Collinsella* as a genus that differed in relative abundance compared to controls[19]; however, the authors noted considerable heterogeneity between studies, suggesting that further investigation is needed to fully elucidate changes in the gut microbiome with epilepsy.

*Collinsella* has been recognized as a pro-inflammatory genus that has been found as a risk factor in several additional neurological diseases of humans, including Alzheimer’s disease[16] and Parkinson’s disease[32]. Many studies[16, 32, 50] have shown positive association between *Collinsella* and the expression of IL-17A, a pro-inflammatory cytokine. This cytokine can cross the blood-brain barrier (BBB) and affect the central nervous system (CNS) by enhancing neuroinflammation, which could lead to the development of epilepsy[51–53].

One of the major products of microbial metabolism in the gut, through alcohol fermentation and the breakdown of non-digestible carbohydrates, is short-chain fatty acids (SCFAs), which have primarily an anti-inflammatory effect[54]. SCFAs are proposed to mitigate gut inflammation and neuroinflammation by multiple mechanisms, including enhancing the BBB, regulating neurotransmission, and affecting microglia[55]. A study of pregnant women showed that the abundance of *Collinsella* is negatively correlated with their fiber intake, with low dietary fiber intake inhibiting the growth of bacteria that produce SCFAs[56]. Furthermore, a deficiency of SCFA producing bacteria was noted in fecal samples from epileptic dogs compared to healthy controls, supporting a role for SCFAs in the development of epilepsy[20]. Besides *Collinsella*, many of the top candidates from our differential abundance analysis are known to alter SCFA production, such as members in *Ruminococcus* and *Clostridium* genera*, Faecalibacterium* genus, and bacteria from the *Lachnospiraceae* family[57, 58]. However, the role of each bacterium could be complex. For example, an ASV from the species *Ruminococcus gnavus* was found significantly more abundant in IE dogs. Although this genus contributes to SCFA production, its abundance is also positively correlated with inflammation[58, 59]. Furthermore, functions of microbiome species can be different at strain level[60, 61]. In the future, direct measurements of SCFAs will improve our ability to investigate this potential causal pathway in canine epilepsy and other neurological diseases.

*Lactobacillus* has been hypothesized to play a role in epilepsy as it can produce gamma aminobutyric acid (GABA), which is an inhibitory neurotransmitter that may indirectly influence the CNS. A study in mice suggests that a GABA-producing *Lactobacillus* strain may have effects on CNS and thereby modulate stress and depression-like behavior[62]. In this study, we observed a significantly higher abundance of one *Lactobacillus* ASV in IE dogs. However, this finding can be confounded by sex as 10 out of 11 dogs with this ASV were female., in both of our pilot study^16^ and the study by Garcia-Belenguer et al.^15^, no significant differences in *Lactobacillus* species between IE and control groups was identified. In a more recent pilot study by Garcia-Belenguer et al.^45^, a higher abundance of *Lactobacillus* was found in IE dogs, and the difference disappeared after administering medium chain triglycerides (MCT) diet for dogs in both groups.

We also used random forest predictive modeling to investigate whether there was a whole microbiome association with IE status that was not evident in analyses of individual taxa. While our Random Forest model achieved a prediction accuracy of 0.588, greater than the 0.5 accuracy that would be achieved by random guessing in this balanced cohort, we acknowledge that this level of discriminative power is not high enough to be useful for clinical diagnosis. However, the better-than-random prediction achieved from whole-microbiome data alona is consistent with the existence of microbial signatures associated with IE.

In this study, we employed a pairwise case-control observational study design. Each pair of animals was selected from the same household while they were fed with the same diet. This design avoids the potential effect induced by diet and living environments. Dogs were not matched with respect to age, sex or breed by design, as requiring these factors as inclusion criteria would have severely limited study recruitment. The household effect was controlled for by leveraging the paired study design in later analyses. When identifying DA taxa, an ensemble DA approach was used to reduce false-positive findings, and we analyzed the ASVs from the one flagged by most methods. However, there were some limitations in our study. Firstly, the ensemble DA method reported the most robustly significant differentially abundant ASVs, but other ASVs highlighted by fewer methods could also be meaningful.

Secondly, relying on 16S rRNA gene sequencing has limited taxonomic resolution and provides no direct functional information, confining our conclusions to the association level. While we proposed possible pathways of how microbiomes could contribute to canine epilepsy, the lack of multi-omics data prevented a deeper exploration of the specific genes or metabolites. Further studies and validations are required to establish the causal relationships between these microbial signatures and the disease.

## Conclusion

This study compares the gut microbiome between IE and control dogs. The abundance of *Collinsella*, a bacterium previously associated with inflammation, was found to differ between the groups. Household was found to be the strongest factor influencing the canine gut microbiome, with microbial compositions being substantially more similar in dogs from the same household than from a different one. Given the frequency of multi-dog ownership, we suggest that other studies of the canine microbiome may also benefit from a similar paired-by-household study design as we used here. Phenobarbital is an effective method to control seizure, and it does not change the large-scale patterns of the microbial communities. Besides IL-17A and SCFAs, future perspectives could be measuring other markers of inflammation in the gut environments and investigate their role in canine epilepsy. Future studies can also evaluate the microbiome in IE dogs over time using longitudinal study design, with focus on *Collinsella* and seizure control. The effect of long-term supplementation of probiotic or fiber diet on the microbiome in IE dogs could also be worthwhile to study in the future.

## Methods

### Study design

#### Dog selection

Dogs were enrolled in the study as pairs, consisting of a dog with IE and a control dog without IE from the same household. To be included in the epilepsy group, dogs were required to have a diagnosis of idiopathic epilepsy based on a Tier 1 confidence level as proposed by the International Veterinary Epilepsy Task Force[63]. In addition, dogs were required to not have received any ASM for a minimum of 30 days prior to study participation (drug-naïve group), or be administered only phenobarbital for at least 3 months and have achieved a minimum serum concentration of 15 µg/ml (drug-treated group). Furthermore, both dogs of the pair were required to: 1) not be receiving any medications aside from parasite preventatives (and phenobarbital if in the drug-treated group); 2) have stool that was considered normal for that dog (as determined by the owner) for at least 2 weeks prior to enrollment; 3) have no evidence of intestinal parasites on fecal examination; and 4) be fed the same diet. Dog pairs were not matched with respect to age, sex, or breed. Owners were required to provide informed consent prior to their dogs’ participation in the study. The study was approved by the NC State University Institutional Animal Care and Use Committee (18-133-0) and the Institutional Review Board (16729).

#### Fecal sample collection

For dogs that met the inclusion criteria, owners were provided collection and shipping materials, including detailed, written instructions on sample collection, storage and shipment. Owners were instructed to collect a freshly defecated fecal sample free of ground contamination from both dogs, putting a portion of each sample in a specimen container. The specimen container was refrigerated until shipped to the investigators via overnight priority service. Once received, a portion of the refrigerated sample was processed for fecal flotation to examine for parasites, and the remainder of the samples kept frozen at -80°C until analyzed.

##### Owner questionnaire

Owners were instructed to complete an online questionnaire using Qualtrics survey software at the time of sample submission. The questionnaire was developed from those utilized by the authors in previous epilepsy studies, and included information on demographics, diet, environment, age of onset of seizures, epilepsy treatment history, seizure frequency over the past 6 months, and history of cluster seizures or status epilepticus. Based on the owners’ responses regarding seizure frequency, IE dogs were grouped according to whether or not they were free from seizures over the last 6 months, and seizure control was classified as being satisfactory (< 1 seizure per month) or unsatisfactory (1 or more seizure per month).

#### Fecal analysis

Fecal samples underwent routine screening for intestinal parasites using standard float protocols and light microscopy. Briefly, 2 grams of the refrigerated fecal sample was thoroughly mixed with 10 mL of FecaMed sodium nitrate solution (Vedco, Inc, St. Joseph, MO) using a wooden popsicle stick. The mixture was filtered through 1 layer of gauze and centrifuged for 5 min at 1,300 rpm. After settling for 10 min, the top layer was removed, placed on a slide, coverslipped, and evaluated under the microscope for the presence of parasite eggs. If a sample screened positive, the dog pair was excluded from further analysis. Owners of such dog pairs were advised to treat the intestinal parasites and collect another fecal sample from both dogs a minimum of 4 weeks after treatment. Resubmitted samples were again screened for intestinal parasites, and samples determined to be clear were included in the study.

### Sequencing

#### DNA Isolation

Fecal samples were transferred to a 2 mL tube containing 200 mg of 106/500μm glass beads (Sigma, St.Louis, MO) and 0.5 ml of Qiagen PM1 buffer (Valencia, CA). Mechanical lysis was performed for 40 minutes on a Digital Vortex Mixer. After a 5-minute centrifugation, 0.45 ml of supernatants was aspirated and transferred to a new tube containing 0.15 ml of Qiagen IRS solution. The suspension was incubated at 4°C overnight. After a 5-minute centrifugation, supernatant was aspirated and transferred to deep well plates containing 0.45 ml of Qiagen binding buffer supplemented with Qiagen ClearMag Beads. DNA was purified using the automated KingFisher™ Flex Purification System and eluted in DNase free water[64–66].

#### 16S rRNA gene amplicon sequencing

12.5 ng of total DNA were amplified using universal primers targeting the V4 region of the bacterial 16S rRNA gene[64, 65]. Primer sequences contained overhang adapters appended to the 5’ end of each primer for compatibility with Illumina sequencing platform. The primers used were F515/R806[67]. Master mixes contained 12.5 ng of total DNA, 0.5 µM of each primer and 2x KAPA HiFi HotStart ReadyMix (KAPA Biosystems, Wilmington, MA). The thermal profile for the amplification of each sample had an initial denaturing step at 95°C for 3 minutes, followed by a cycling of denaturing of 95°C for 30 seconds, annealing at 55°C for 30 seconds and a 30 second extension at 72°C (25 cycles), a 5-minute extension at 72°C and a final hold at 4°C. Each 16S amplicon was purified using the AMPure XP reagent (Beckman Coulter, Indianapolis, IN). In the next step, each sample was amplified using a limited cycle PCR program, adding Illumina sequencing adapters and dual index barcodes index 1 (i7) and index 2 (i5) (Illumina, San Diego, CA) to the amplicon target. The thermal profile for the amplification of each sample had an initial denaturing step at 95°C for 3 minutes, followed by a denaturing cycle of 95°C for 30 seconds, annealing at 55°C for 30 seconds and a 30 second extension at 72°C (8 cycles), a 5-minute extension at 72°C and a final hold at 4°C. The final libraries were again purified using the AMPure XP reagent (Beckman Coulter), quantified and normalized prior to pooling. The DNA library pool was then denatured with NaOH, diluted with hybridization buffer, and heat denatured before loading on the MiSeq reagent cartridge (Illumina) and on the MiSeq instrument (Illumina). Automated cluster generation and paired–end sequencing with dual reads were performed according to the manufacturer’s instructions.

### Data Curation

The DADA2 R package[68] (version 1.33.0) was used to determine the microbiome composition from the paired-end fastq files. Using the filterAndTrim function, read pairs were truncated to forward and reverse lengths of 200 and 150 nucleotides, and primers were removed by cutting out the first 19 and 20 nucleotides, respectively. The dada function was used to denoise the trimmed and filtered reads, and the “pseudo-pooling” mode was used to increase the sensitivity of the DADA2 algorithm on rare variants. The removeBimeraDenovo function was used to detect and remove the chimeric sequences. Taxonomy was first assigned to genus-level using the naïve Bayesian classifier method in assignTaxonomy function with the 16S Silva reference database[69] (version 138.1) and then assigned to species-level by exact matching to the Silva reference database using the addSpecies function.

A phylogenetic tree was constructed using the DECIPHER R package[70] (version 3.2.0). ASV sequences were first aligned using the AlignSeqs function. Multiple sequence alignments were used in the Treeline function with the minimum evolution method.

The phyloseq R package[71] (version 1.50.0) was used to store and manage data. The metadata, sequence table, taxonomy table, and phylogenetic tree were stored in a phyloseq object as the non-rarefied dataset. To maintain the pairwise structure of the study, the total set of samples was reduced so that each household included just one IE dog sample and one control dog sample. For duplicated samples, the one with the highest read depth was kept.

A rarefied dataset was created using the rarefy_even_depth function from the phyloseq R package to control the effect of uneven sequencing depth. Two samples were found to have low read depth, and so those two households were removed from the study. The rarefaction threshold was set to 44590 to maximize the read depth while minimizing the loss of sample size. The rarefied dataset was used in all analyses except for the DA analysis.

In order to compare the phylum-level taxonomic profiles of our current study to our pilot study, taxonomy was re-assigned to the pilot study ASVs using the Silva reference database (version 138.1), and the phylogenetic tree was constructed using the DECIPHER R package. One pair of samples was removed due to the low read depth. The pilot study sequence table and metadata can be downloaded from https://github.com/benjjneb/CanineEpilepsyManuscript. The pilot study data was not included in any of the other analyses reported here.

### Statistical Analysis

Pearson’s Chi-square test was used to identify if the IE status of dogs was associated with breed group or sex. The association between IE status and age was evaluated by t-test. The difference of large-scale microbial patterns between IE and control dogs and other covariates were analyzed with Permutational Multivariate Analysis of Variance (PERMANOVA) tests based on Bray-Curtis and Weighted-Unifrac distances using the adonis2 function from the VEGAN R package[72] (version 2.6-8). To avoid the confounding effect introduced by households, when analyzing the microbiome difference between IE and control dogs, 1000 permutations were calculated with the group label exchange restricted in each household. When analyzing the effect of breed group, age, sex, and phenobarbital administration, the household effect was first taken into account to control its potential confounding effect using the model:

#### Dissimilarity∼Household+Factor

Only IE dogs were considered when analyzing the effect of phenobarbital administration. The Shannon index was calculated by the estimate_richness function from the phyloseq R package and the ANOVA tests were applied to analyze the effect of each factor on alpha diversity using the anova_test function from the rstatix R package[73] (version 0.7.2). Both PERMANOVA and ANOVA tests used Type I sum of squares.

Pairwise_sign_test from the rstatix R package was used to compare the abundance difference of *Collinsella* and *Lactobacillus* ASVs between IE and control dogs. Pearson’s Chi-square test was used to evaluate the association between the presence of *Collinsella* or *Lactobacillus* ASVs and sex. Linear model regression was used to identify if the abundance of *Collinsella* or *Lactobacillus* ASVs will change by age. The abundance difference of *Collinsella* or *Lactobacillus* ASVs between seizure control or seizure freedom groups were calculated by t-test. A significance level of p < 0.05 was established for all analyses. All the analyses in R markdown format can be found in the GitHub repository: https://github.com/Yixuan39/CanineEpilepsy2

### Differential Abundance Analysis

We used multiple DA tools to find ASVs that are reported as differentially abundant between IE and control groups across tools. To reduce the number of multiple tests, we set a threshold that excluded ASVs with prevalence lower than 10%. Under this threshold, we decreased the number of taxa from 2879 to 387 and retained around 96% of reads on average. We evaluated the results from each tool by comparing the returned p-value and estimated effect size. Adjusted p-value for multiple comparisons was not used in this study. Different tools might use different measurements for the effect size. Here we unified the effect size as the LFC by transforming the effect size values from different tools. The LFC values of corncob were not reported since the software only estimates relative abundance differences instead of LFC. However, the direction of estimated relative abundance differences can be compared to the LFC values calculated by other tools.

#### ALDEx2

We used the non-rarefied count data with the aldex.ttest and aldex.glm function from the ALDEx2 R package (version 1.38.0). 128 Monte-Carlo instances of Dirichlet distributions were generated for each sample, and the count data were centered log2-ratio (CLR) transformed. Then, we performed pairwise Welch’s t-test between the IE and control groups and reported the posterior predictive p-value of Welch’s t-test and the LFC between the two groups. We built an ALDEx2 generalized linear model with the health condition group and household added as factors and reported the p-value of the term of the health condition group with the estimated centered log2-ratio difference.

#### ANCOM-BC

We used the non-rarefied count data with the ancombc function from the ANCOMBC R package (version 2.8.0). We built a negative binomial regression model with the health condition group and household added as factors and reported the p-value of the term of the health condition group with the estimated log2 fold change converted from natural log fold change.

#### corncob

We used the non-rarefied count data with the differentialTest function from the corncob R package (version 0.4.1). The function fitted each taxon based on the beta-binomial distribution for the null and alternative models. The null model was set with only the household factor and the alternative model was set with the healthcondition group and the household factor. Then, likelihood ratio tests were performed, and the p-value was reported.

#### LinDA

We used the non-rarefied count data with the linda function from the MicrobiomeStat R package (version 1.2). The function normalized the raw read counts with CLR transformation and fitted each taxon with the linear mixed-effect regression models. We added the health condition group and household as factors to the modeland reported the p-value of the term of the health condition group with the estimated log2 fold change.

#### Pairwise T-test

We used the rarefied count data with the t.test function from the basic R software (version 4.4.1). We performed the pairwise t-test for each taxon between the IE and control groups and reported the p-values. The LFC values were calculated as the log2 ratio of mean relative abundances, employing a pseudo-count of 0.5 to facilitate the inclusion of taxa with zero counts in either group.

### Evaluating ASV sharing between dogs from the same household

When comparing the similarities between dogs from different households, we created a new dataset that each IE dog was paired with a control dog from a different household. As 33 out of 49 households had dogs with the same predominant breed based on AKC breed guidelines, we wanted to make sure the same proportion of pairs in the new dataset had the same predominant breed to control the breed effect. To do this, when selecting dogs, we randomly picked an unaffected dog with the same predominant breed when possible. With this strategy, 31 pairs of same predominant breed dogs from different households were created. To balance the breed effect, two of these paired dogs were randomly removed to keep both datasets containing 31 out of 47 households that had dogs with the same predominant breed.

Pairwise similarities for both datasets based on Jaccard distances were measured using the distance function from the phyloseq R package. Pairwise T-tests were used to compare the similarities for different community-wise distances between dogs from the same households and dogs from different households To determine if the presence of ASVs within each household was independent, the observed proportion of households in which a given ASV was present in both dogs (P_Observed_) was compared to the null expectation of random co-occurrence (P_null_). Given the sample size of the study is 98, (P_null_) was calculated as 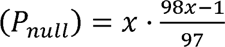, where x represents the study-wide prevalence of the ASV.

### Machine Learning Prediction

#### Data curation

The dataset used for machine learning prediction included the dog’s health status as the response variable, the household as the grouping variable for sampling, and the microbial composition in the form of per-ASV relative abundances as predictors. For the microbial data, only ASVs with prevalence greater than 10% were used, and the compositions were transformed into proportions.

#### Dimension reduction

Two dimension reduction methods were used in this study. The first method converted the microbial data from ASV-level to Phylum-level using the tax_glom function from the phyloseq R package (version 1.50.0). The second method converted the microbial data from ASV-level to orthogonal axes using PCoA technique. Axes were calculated using the ordinate function from the phyloseq R package (version 1.50.0) with the PCoA method based on Bray-Curtis distance. The first 30 axes were selected as predictors in the model.

#### Model training

The Random Forest model from the ranger R package[74] (version 0.17.0) was employed in this study. With the rsample R package[75] (version 1.2.1), the dataset was first split into training (70%) and testing (30%) parts by household using the group_initial_split function. Then, 10-fold cross validation was performed during the model training process using the group_vfold_cv function. The hyperparameters were tuned by Bayesian Optimization using the tune_bayes function from the tune R package[76] (version 1.2.1). The maximum number of iteration searches of the tuning process was set to 40, and the searching process will be terminated if the results have no improvement in 10 iterations. As the results were strongly influenced by the initial data splitting, for each dataset, we calculated the model accuracy 30 times. All the functions used in the training and testing steps were included in the tidymodels ecosystem (version 1.2.0).

## Declarations

### Ethics approval and consent to participate

This study was approved by North Carolina State Institutional Animal Care and Use Committee (Protocol 18-133-O) and Institutional Review Board (Protocol 16729).

Owners of dogs were required to provide informed consent prior to participation in the study.

## Consent for publication

Not applicable.

## Availability of Data and Material

The raw 16S rRNA gene sequencing data has been deposited in the SRA under the deposition PRJNA1308174. The computational analyses performed in this study are available in R markdown format at the GitHub repository: https://github.com/Yixuan39/CanineEpilepsy2

## Authors’ Contributions

KM conceived and designed the study. KM and BC obtained funding. KM and JN recruited study participants and collected samples. MAP performed the 16S rRNA gene sequencing. YY performed data analysis and interpretation under supervision by BC. YY wrote the manuscript, with contributions from KM, JN and BC. All authors reviewed and approved the final manuscript.

## Acknowledgements

We acknowledge the dogs and their owners who generously participated in this study.

